# More than meets the eye: syntopic and morphologically similar mangrove killifish species show different mating systems and patterns of genetic structure along the Brazilian coast

**DOI:** 10.1101/2020.06.30.179937

**Authors:** Waldir M. Berbel-Filho, Andrey Tatarenkov, Helder M. V. Espirito-Santo, Mateus G. Lira, Carlos Garcia de Leaniz, Sergio M. Q. Lima, Sofia Consuegra

## Abstract

Different mating systems can strongly affect the extent of genetic diversity and population structure among species. Given the increased effects of genetic drift on reduced population size, theory predicts that species undergoing self-fertilization should have greater population structure than outcrossed species, however demographic dynamics may affect this scenario. The mangrove killifish clade is composed of the two only known examples of self-fertilising species among vertebrates *(Kryptolebias marmoratus* and *K. hermaphroditus*). A third species in this clade, *K. ocellatus,* inhabits mangrove forests in southeast Brazil, however its mating system and patterns of genetic structure have been rarely explored. Here, we examined the genetic structure and phylogeographic patterns of *K. ocellatus* along its distribution, using mitochondrial DNA and microsatellites to compare its patterns of genetic structure with the predominantly selfing and often syntopic, *K. hermaphroditus*. Our results indicate that *K. ocellatus* reproduces mainly by outcrossing across much of its known range, with no current evidence of selfing, despite being an androdioecious species. Our results also reveal a stronger population subdivision in *K. ocellatus* compared to *K. hermaphroditus*, contrary to the theoretical predictions based on reproductive biology of the two species Our findings indicate that, although morphologically similar, *K. ocellatus* and *K. hermaphroditus* had remarkably different evolutionary histories when colonising the same mangrove areas in south-eastern Brazil, with other factors (e. g. time of colonisation, dispersal/establishment capacity) having more profound effects on the current population structuring of those species than differences in mating systems.

## Introduction

Differences in mating systems can have profound effects on the extent of genetic variation and of population structure (Charlesworth and Wright, 2001). Theory predicts that selfing species should have deeper population structure than outcrossed species, given the stronger effects of genetic drift on reduced population size (Charlesworth, 2003; Meunier *et al*, 2004). At a broader geographic scale, multiple geographically isolated selfing lineages should result in high levels of genetic diversity in selfing species (Avise and Tatarenkov, 2015). However, this pattern can be influenced by temporal population dynamics, such as dispersal and colonisation capacity (Siol *et al*, 2007).

Although the impact of different mating systems on population structure has already been explored in plants (Willi and Määttänen 2011), this research has lagged behind in animal systems, particularly on vertebrates, where most species are dioicous and are obligate outcrossing (Jarne and Auld 2006). However, a unique diversity of mating systems exists in the mangrove killifish species of the genus *Kryptolebias* (Costa *et al*, 2010; Avise and Tatarenkov 2015). The mangrove killifishes clade is composed of the only representatives among all rivulids (350+ species) living in brackish waters (Costa *et al*, 2010), and the only two known examples of self-fertilising hermaphroditism among vertebrates (*K. marmoratus* and *K*. *hermaphroditus,* species that form the “*K*. *marmoratus* species complex”, see Tatarenkov *et al* (2017)). A third species in the mangrove killifish clade is *Kryptolebias ocellatus* (previously known as *Kryptolebias caudomarginatus,* taxonomic nomenclature still under discussion (Costa, 2011; Huber, 2017)). *Kryptolebias ocellatus* is endemic to intermittent mangrove microhabitats in southern and south-eastern Brazil (Costa, 2016).

*Kryptolebias ocellatus* has been historically bred in aquaria, and while behavioural observations indicate it reproduces via outcrossing (Seegers, 1984), its populations are composed of males and simultaneous hermaphrodites (Costa *et al*, 2010), leaving open the possibility that this species may also undergoes self-fertilisation. However, while the genetic analysis of two populations in this species found no evidence for selfing (Tatarenkov *et al*, 2009), the possibility of self-fertilisation at a broader geographical scale cannot be ruled out, as rates of selfing and outcrossing are known to vary geographically in the mixed-mating *Kryptolebias* species (Berbel-Filho et al, 2019; Tatarenkov et al. 2011). In the northernmost part of its distribution (Guanabara and Sepetiba Bays, 22° S), *K*. *ocellatus* is often syntopic (i.e. coexisting at the same habitat at the same time) with *K. hermaphroditus* (Costa, 2011; Costa, 2016), a species composed mostly of self-fertilising hermaphrodites and very rare males (Berbel-Filho *et al*, 2016; Costa, 2016), resulting in occasional outcrossing but at very low frequencies (Berbel-Filho *et al*, 2019; Tatarenkov *et al*, 2017). Extremely low levels of genetic diversity in *K*. *hermaphroditus*, especially at the southernmost edge of its distribution (where it is syntopic with *K*. *ocellatus*), suggest relatively recent dispersal and colonisation of this species in south-eastern Brazil (Tatarenkov *et al*, 2009; 2011; 2017).

*Kryptolebias ocellatus* and *K. hermaphroditus* coexist in shallow mangrove microhabitats, such as. temporary pools and crab burrows in discontinuous patches of mangrove forests in south-eastern Brazil and display very similar body shape and colour patterns (Costa, 2016; Tatarenkov *et al*, 2017) (Fig. 1). For these reasons, morphologically-based taxonomic classification of the two species has been historically difficult (Costa, 2006; Costa, 2011; Costa, 2016; Huber, 2017). However, phylogenetic studies indicate that *K. ocellatus* is the sister-species of the clade containing the two selfing species from *‘K. marmoratus species complex’ (K. marmoratus* and *K*. *hermaphroditus)* (Kanamori *et al*, 2016; Tatarenkov *et al*, 2009; Vermeulen and Hrbek, 2005), suggesting that the current syntopy between congeners (*K. hermaphroditus* and *K*. *ocellatus*) in south-eastern Brazil is more likely due to dispersal and colonisation rather than to local speciation.

**Figure 1.**
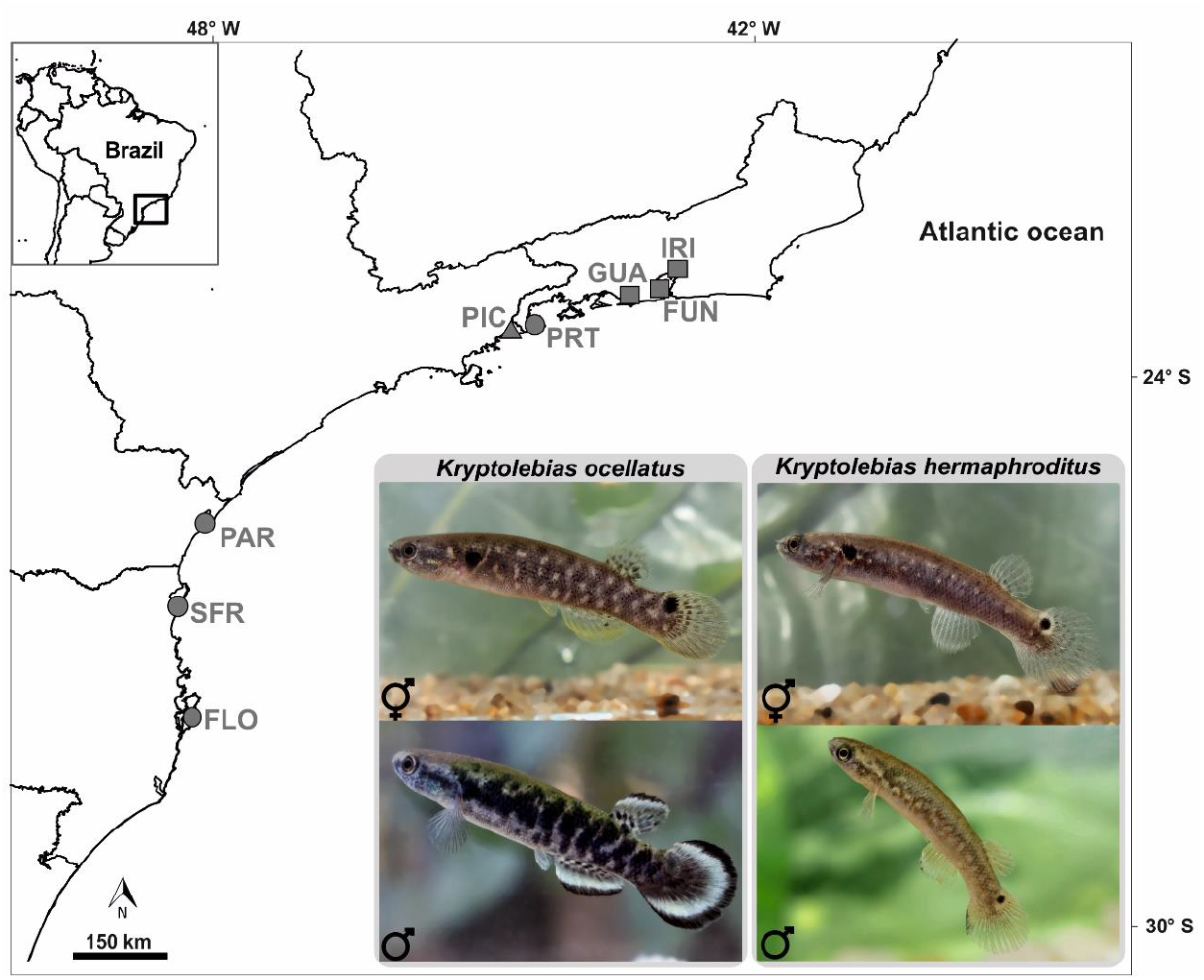
Sampling locations for *Kryptolebias ocellatus.* Squares represent locations where *K. ocellatus* and *K. hermaphroditus* are syntopic, circles are for locations where only *K. ocellatus* is found, while triangle designates site where only *K. hermaphroditus* is found. Labels for locations are described on Table 1.

Here, we investigate the population structure of *K*. *ocellatus* across its range using mitochondrial DNA and microsatellite markers. To test the potential role of different mating systems in determining the population structure, we compared the patterns of genetic structure and diversity of *K*. *ocellatus* with previously-published data for the self-fertilising species *K*. *hermaphroditus*.

## Material and Methods

### Sampling collection

We sampled *K. ocellatus* in southern and south-eastern Brazil, covering most of its known range (Costa, 2016), between August and September 2017 (Fig. 1). Mangrove forests along this ~900 km long coastal area is discontinuous and heavily fragmented by urbanisation (Barletta and Lima, 2019; Branoff, 2017). We collected the fish using hand nets in mangrove temporary pools and crab burrows (Fig. 1; Table 1). Sex (male or hermaphrodite) was inferred by body and fin coloration patterns, which are reliably used for sex differentiation in mangrove killifish species (Scarsella *et al*, 2018). In *K. ocellatus,* males were identified by a black spot on the dorsal part of the caudal fin (Costa, 2016). In *K. hermaphroditus,* males were identified by the presence of by broad black margin along the whole caudal fin, bordered by a broad sub-marginal white zone as described in Costa (2016).

**Table 1.**
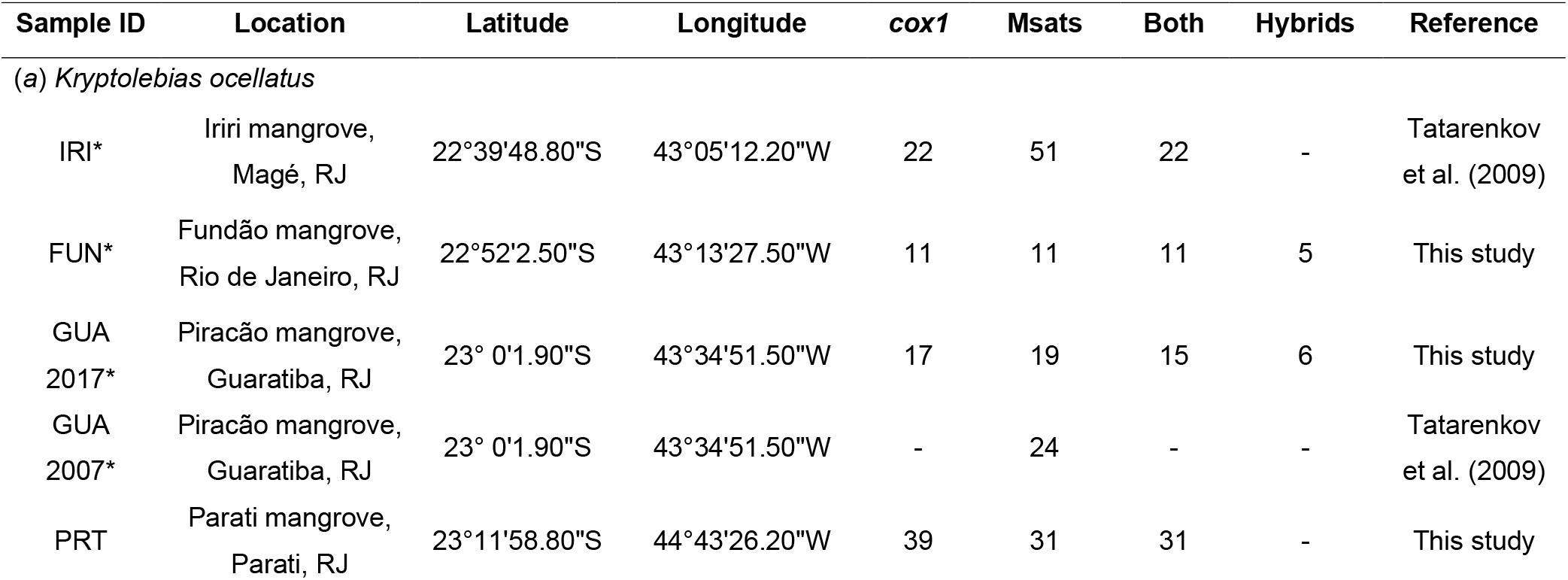
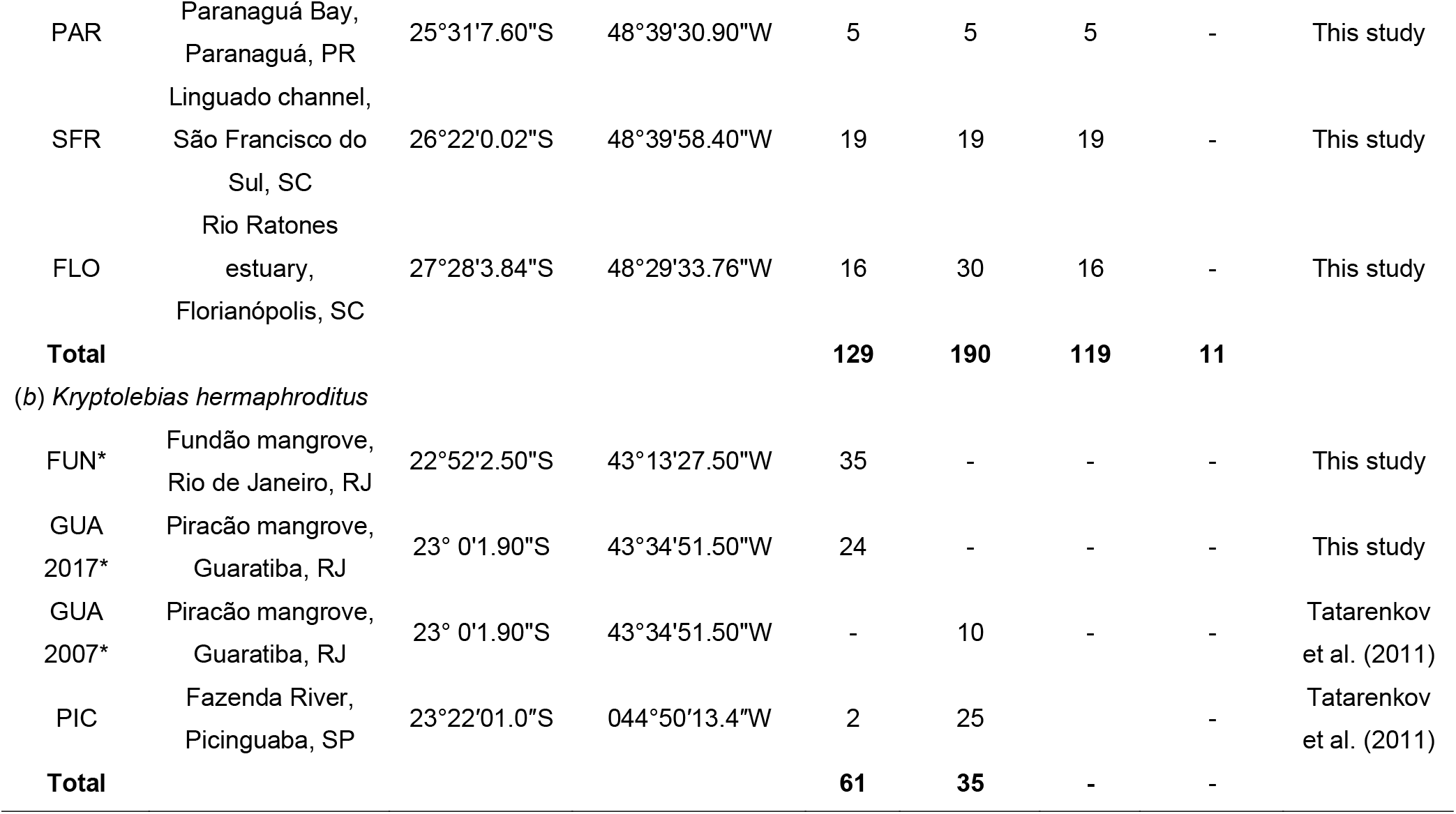
Sampling locations and sampling sizes in (*a*) *Kryptolebias ocellatus* and (*b*) *Kryptolebias hermaphroditus.* PR, Paraná State; RJ, Rio de Janeiro State; SC, Santa Catarina State; SP, São Paulo State. Asterisks denotes sampling locations where populations are syntopic. “Both” refers to the number of individuals sampled for both mtDNA and genotyped for 16 microsatellites. “Hybrids” refers to individuals with evidence of admixed genetic backgrounds between *K. ocellatus* and *K. hermaphroditus.* “Reference” refers to the source of microsatellite data.

### Genetic markers

A subset of 16 microsatellites from Mackiewicz *et al* (2006) was amplified and genotyped following Tatarenkov *et al* (2010). The mitochondrial gene cytochrome oxidase subunit I *(cox1)* was also used to investigate the genetic structure and major mtDNA lineages distribution.

A 618 bp region of the *cox1* was amplified with FishCOI-F (5’-TCAACYAATCAYAAAGACATYGGCAC-3’) and FishCOI-R (5’-ACTTCYGGGTGTCCRAARAAYCA-3’) primers as in Tatarenkov *et al* (2017). Both forward and reverse DNA strands were Sanger sequenced and assembled using Geneious v. 9.1.8 (www.geneious.com). Sequences were deposited in GenBank (accession numbers: *K. ocellatus*: MN400774 – MN400902; *K. hermaphroditus*: MN400903 – MN400963).

### mtDNA and microsatellites datasets

We combined newly generated sequences and genotypes with data from previous studies (Tatarenkov *et al*, 2011; Tatarenkov *et al*, 2009) for the present genetic analyses (an update of the current taxonomic nomenclature for the study species, which changed in the last years, is provided in Supplementary material).

The *K. ocellatus* dataset consisted of individuals from seven sampling locations, three of them (IRI, FUN and GUA in Fig.1) where the species was found in syntopy with *K. hermaphroditus* in southeast Brazil, and four (PRT, PAR, SFR and FLO in Fig. 1), where only *K. ocellatus* is found. Overall, 200 *K. ocellatus* individuals were analysed, 119 (59.5%) of them were both sequenced for *cox1* and genotyped for 16 microsatellites (Table 1). In addition, 10 individuals were sequenced only for *cox1* and 71 individuals were only genotyped for microsatellites (Table 1), resulting in 129 individuals sequenced for *cox1* and 190 individuals genotyped for microsatellites (Table 1).

In the case of Iriri population (IRI in Fig. 1), new *cox1 K. ocellatus* sequences were obtained for 22 of the 51 individuals previously genotyped for microsatellites in Tatarenkov *et al*(2009) (Table 1). The *K. ocellatus* microsatellite dataset for Guaratiba (GUA in Fig. 1) consisted of 19 individuals sampled in 2017 (17 of them with *cox1* data) and 24 genotypes from individuals sampled in 2007 (no *cox1* data) reported in Tatarenkov *et al* (2009) (Table 1).

To compare the patterns of genetic structure and diversity between *K. ocellatus* and *K. hermaphroditus* in south-eastern Brazil, we generated a *cox1* dataset for *K. hermaphroditus* consisting of 61 sequences from three locations in southeast Brazil (FUN, GUA and PIC in Fig. 1), two of them (FUN and GUA in Fig. 1) representing areas of syntopy for both species (Fig 1; Table 1). We also used 35 *K. hermaphroditus* microsatellites genotypes from Tatarenkov et al. (2011), comparing both species for 14 of the 16 microsatellites amplified here for *K. ocellatus* (R34 and R112 were not genotyped in *K. hermaphroditus*). *Kryptolebias hermaphroditus* individuals had been sampled from two populations in south-eastern Brazil (PIC and GUA in Fig. 1) in 2007.

### mtDNA phylogenetic and phylogeographic analyses

A Bayesian coalescent reconstruction was carried out using BEAST v. 2.5.1 (Bouckaert *et al*, 2014). The sequences included the *cox1* haplotypes found across129 *K. ocellatus* individuals, as well as the following outgroups: the single haplotype found across 61 *K. hermaphroditus* individuals (see results) in south-eastern Brazil; two sequences of *K. marmoratus* (accession numbers: MF555022.1 and MF554974.1) and two sequences of the ‘Central clade’ lineage (accession numbers: MF555047.1 and MF555072.1), a selfing lineage present in Central America and Caribbean. The ‘Central clade’ is closely related to *K. hermaphroditus* (Tatarenkov et al. 2017); however, its formal taxonomic status is still under debate. The best-fit model of nucleotide substitution was selected according to the Akaike and Bayesian Criteria on jModelTest2 (Darriba *et al*, 2012). The substitution model indicated by jModelTest2was the 3-paratemer model with unequal base frequencies and invariant sites (TPM1uf+I). To time-calibrate the phylogenetic reconstruction and allow for rate variation among lineages, a lognormal relaxed molecular clock of 0.009 substitutions per site per million years was used, based on the *cox1* Goodeidae fossil-calibrated molecular rate described in Webb *et al* (2004). We performed three independent runs of 10^6^ Markov Chain Monte Carlo (MCMC) steps, sampling every 10^3^ steps. Tracer v. 1.7.1 (Rambaut *et al*, 2018) was used to assess convergence and effective sample sizes (≥ 200) among MCMC runs. The software TREEANNOTATOR v. 2.5.1 (Bouckaert *et al*, 2014) was used to discard the first 200 trees (20%) as burn-in, and to generate a consensus tree with posterior probability value for each clade.

For *K. ocellatus,* the number of haplotypes *(H)* and polymorphic sites (S), haplotype (*h*) and nucleotide diversities (π) for each sampling location and major mtDNA clades were calculated using DNAsp v. 6.10.04 (Rozas *et al*, 2017). For generating pairwise fixation indices (F_ST_) among major clades and sampling locations, we used Arlequin v. 3.5.2.2 (Excoffier and Lischer, 2010). We used Mega v. 7.0.26 (Kumar *et al*, 2016) to calculate Kimura-2-Parameter (K2P) genetic distances among major clades (see results) and sampled populations. To visualise haplotypes distribution and divergence, we reconstructed a *cox1* haplotype network using POPART (Leigh and Bryant, 2015).

### Genetic structuring and clustering analysis based on microsatellite data

For the microsatellite data, Micro-checker v. 2.2 (van Oosterhout *et al*, 2004) was used to check for errors in the data and presence of null alleles. To assess overall differentiation at the population level, we used FSTAT v. 2.9.3.2 (Goudet, 1995) to calculate F_ST_ and conduct exact G-tests based on 10,000 randomizations of alleles. FSTAT was also used to measure departures from Hardy–Weinberg equilibrium. *P* values for F_IS_ for each locus were based on 2240 randomizations, and *P* values over all loci were calculated from a weighted average of the statistic obtained for each locus. Unbiased expected (H_E_) and observed heterozygosity (H_O_) were calculated using MSA v. 4.05 (Dieringer and Schlötterer, 2003).

We generated a Neighbor-joining tree with 1000 bootstrap replications using Poptree2 (Takezaki *et al*, 2010) based on a matrix of pairwise Nei’s genetic distances between sampling points. The overall genotypic associations of individuals were visualized with a factorial correspondence analysis (FCA) using the procedure implemented in GENETIX v. 4.04 (Belkhir, 2004).

We applied three different methods to estimate the most likely number of genetic clusters (K) across *K. ocellatus* distribution. First, using only microsatellite data we ran STRUCTURE 2.3.4 (Pritchard *et al*, 2000) with the following parameters: K values ranging 1–10, 10 iterations per K, a total of 1,000,000 MCMC with 100,000 burn-in, admixture model, independent allele frequencies. To identify the uppermost hierarchical level of genetic structure, we chose the most likely K value using second-order rate of change of likelihood ΔK method (Evanno *et al*, 2005), implemented in Structure Harvester (Earl, 2012). Independent STRUCTURE runs were aligned and plotted using CLUMPAK (Kopelman *et al*, 2015).

Given the uneven number of individuals in our sample, we also used STRUCTURESELECTOR (Li and Liu, 2018), which provides four metrics of cluster estimates o identify the most likely number of genetic clusters (median of means (MedMeaK), maximum of means (MaxMeaK), median of medians (MedMedK) and maximum of medians (MaxMedK)) (Puechmaille 2016).

### Genetic structuring based on mtDNA and microsatellites data

To integrate mtDNA, microsatellites data and spatial information, we used Geneland v. 4.0.8 (Guillot *et al*, 2008), which takes into account spatial information from each individual, also allowing for uncertainty in the positioning of sampled individuals. To identify spatial population distribution and assess individual assignment to the most likely K, we followed Guillot *et al* (2005). As Geneland allows to incorporate individuals with missing data from one of the markers, we combined mtDNA and microsatellites data for a total 200 *K. ocellatus* individuals, including individuals without mtDNA (71 individuals) or microsatellite genotypes (ten individuals). To avoid any bias potentially introduced by introgression on genetic structure patterns, we removed microsatellites data for the eleven potential hybrid individuals (see results), however maintained their mtDNA for Geneland, as they showed no evidence of introgression and would represent the mtDNA of the parental individuals (see results). Therefore, Geneland was run with information from 200 individuals, 129 with mtDNA information, 179 with microsatellites genotypes, and 108 (after removal of microsatellites data from 11 hybrids) with information for both markers. We repeated the analysis excluding the hybrid individuals from the mtDNA to assess their contribution to the results. Geographical coordinates (geo-referenced according to the sampling points and with uncertainty of ±0.05 in both latitude and longitude) were included for all individuals. K ranges from 1 to 10. Ten multiple runs were performed with 10,000,000 MCMC iterations, sampled every 1,000 iterations. Once the most likely K value was inferred from the modal value across the 10 multiple runs, we ran the MCMC again with other 10 multiple runs and K fixed to assigned value. These final 10 runs were postprocessed (with a burn-in of 20%) in order to obtain posterior probabilities of population membership for each individual. All Geneland analyses were performed using “geneland” R package (Guillot *et al*, 2008).

### Isolation by distance in *K. ocellatus*

Given the discontinuous distribution of mangrove forest in south-eastern Brazil, we tested the association between geographical and genetic distance in *K. ocellatus.* We estimated the pairwise geographical distance (straight line in kilometres) among sampling points in R v. 3.5.3. We used IBD v. 1.52 (Bohonak, 2002), running a Mantel test between the matrices of pairwise F_ST_ between sampling points (both for mtDNA and microsatellites) and estimated geographical distance in kilometres.

## Results

### mtDNA phylogenetic and phylogeographic analysis

Twenty-two *cox1* haplotypes (618bp-long) were recovered from 129 *K. ocellatus* individuals sequenced. In contrast, only one *cox1* haplotype was found for *K. hermaphroditus* across 61 individuals (Table S1). Overall, our phylogenetic reconstruction grouped all *K. ocellatus* haplotypes in a monophyletic clade, with a sister-clade composed by the selfing mangrove killifish species, namely *K. hermaphroditus* and *K. marmoratus* (Fig. 2). In *K. ocellatus,* a clear geographical pattern was found by the Bayesian reconstruction tree using *cox1* haplotypes (Fig. 2). Three major lineages were found: a clade composed of haplotypes from sampling locations within Guanabara and Sepetiba’s Bays (IRI, FUN and GUA; hereafter called Northern clade), clustered with a clade containing haplotypes from the opposite side of Sepetiba Bay (PRT; hereafter called Parati clade), although the support for the grouping of Northern and Parati clades was low (PP: 0.75). The third clade was composed of haplotypes from sampling points in southern Brazil (PAR, SFR, FLO; hereafter called the Southern clade). These three major clades were also supported by NJ tree using microsatellites distances and the haplotype network (Fig. S1 and Fig. 2).

**Figure 2.**
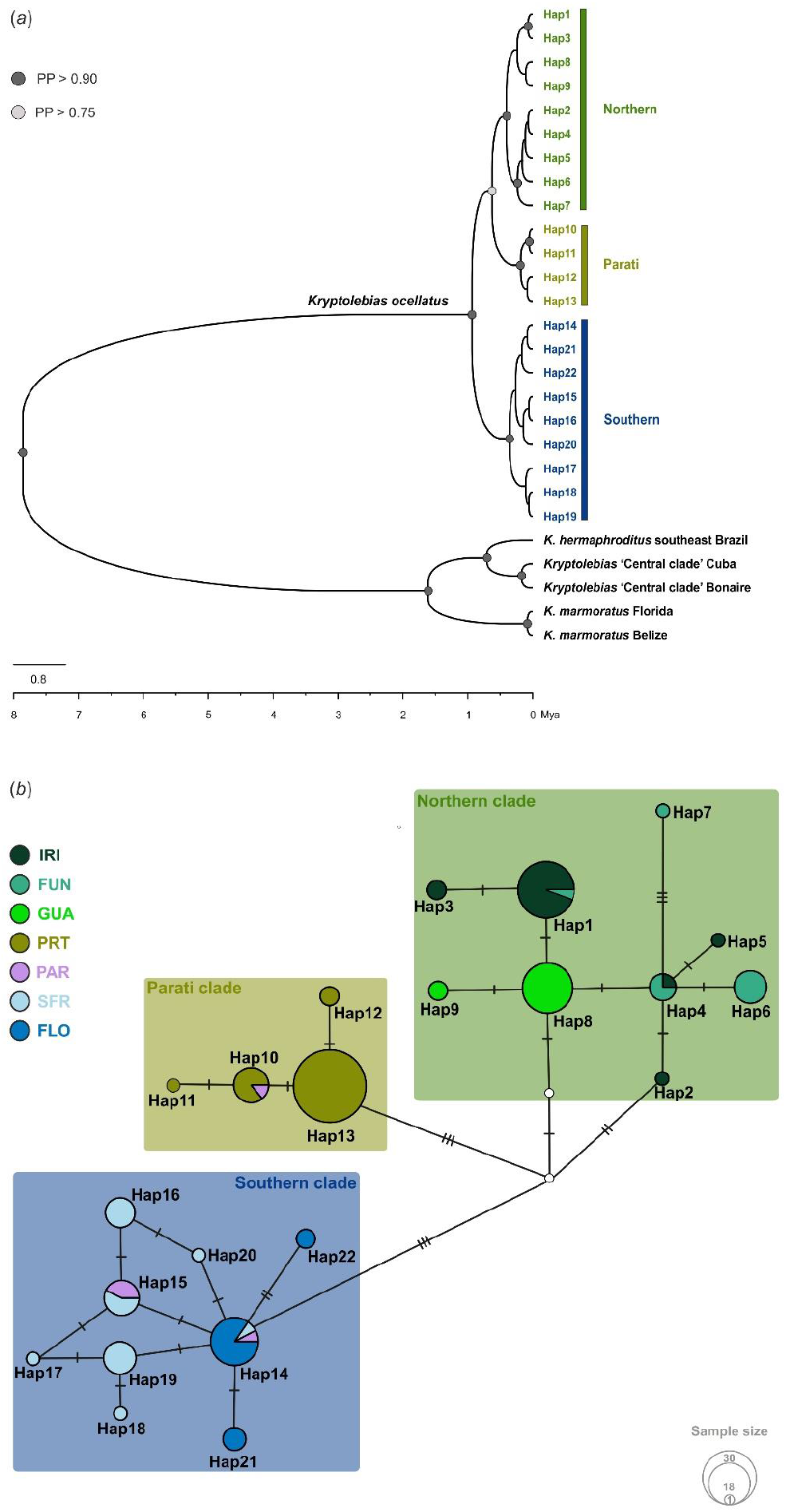
*(a)* Bayesian time-calibrated phylogenetic gene tree for the 22 mitochondrial cytochrome oxidase 1 gene *(cox1)* haplotypes found across129 specimens of *Kryptolebias ocellatus.* The single *cox1* haplotype found for *K. hermaphroditus* in southeast Brazil, two *cox1* haplotypes from ‘Central clade’, which is a lineage related to *K. hermaphroditus* (Tatarenkov et al. 2017) and two *cox1* haplotypes from *K. marmoratus* were used as outgroups. The taxonomic status of the Central clade is in revision (Tatarenkov *et al*, 2020). Circles at nodes represent values of Bayesian posterior probability (PP). Only PP > 0.75 are shown. Scale at the bottom in millions of years (Mya). *(b) cox1* haplotype network for 129 specimens of *K. ocellatus.* Each circle represents a haplotype and its size is proportional to the frequency of the haplotype. Ticks on branches connecting the haplotypes indicate nucleotide mutations. Different colours are used for each locality.

In *K. ocellatus,* overall haplotype diversity was 0.89, being the highest in the SFR (Southern clade) and the lowest in GUA (Northern clade) populations. Nucleotide diversity was generally low (*π*=0.007), and followed a similar pattern to the haplotype diversity, being the highest at PAR (Southern clade) and the lowest at GUA (0.0004). The same pattern of haplotype and nucleotide diversity was seen when sampling locations were grouped according to the major mtDNA clades, with the Southern clade being the most diverse, followed by the Northern and Parati clades, respectively (Table S2).

The average F_ST_ value all pairwise comparisons was 0.72 (sd = ± 0.22) in *K. ocellatus.* All F_ST_ pairwise comparisons among sampling locations were significant, with the exception of the comparison between SFR and PAR (within the Southern clade). The highest F_ST_ value (0.92) was found in the comparison between FLO and PRT (Parati clade), while the lowest (0.14) was found between SFR and PAR (Table S3). All F_ST_ pairwise comparisons were significant when grouping sampling locations into the major mtDNA clades (mean = 0.80). K2P genetic distances followed a similar pattern to the F_ST_ values, with the highest value (1.4%) being observed between samples from the Southern and Parati clades, while the lowest value (0.2%) between samples of the same mtDNA clade (Table S3). K2P genetic distance between *K. ocellatus* and *K. hermaphroditus* was 11.2%.

Additional analysis revealed similar patterns of genetic diversity (Table S3) and genetic differentiation among populations (F_ST_ and genetic distances; Table S5) in a dataset excluding the *cox1* sequences from hybrid individuals (see below), suggesting that the hybridisation has not affected the general patterns of genetic structure observed at the mtDNA.

### Microsatellite variation within populations

We found evidence for possible hybridisation between *K. ocellatus* and *K. hermaphroditus* in two syntopic populations (FUN and GUA; Fig. 3 and Fig. S2). A Structure analysis using both *K. ocellatus* and *K. hermaphroditus* microsatellite data, indicated that the uppermost level of genetic structure was K = 3, with the two genetic clusters within *K. ocellatus* (corresponding to Northern and Southern populations, see below), and a third cluster with *K. hermaphroditus* individuals from GUA and PIC. Eleven individuals had admixed genetic background between *K. ocellatus* and *K. hermaphroditus*, five in FUN and six in GUA (only 2017 sampling) (Fig. 3 and Fig. S2). These potential hybrids exhibited *K. ocellatus* mtDNA haplotypes (haplotypes 4, 6, 7 and 8; Fig. 2; Table S1), all of them (except haplotype 7) commonly found on non-admixed individuals from other northern populations. To avoid any bias caused by these potential hybrids, we excluded them from all population genetics analyses based on microsatellites data. The implication of this hybridisation case will be explored elsewhere.

**Figure 3.**
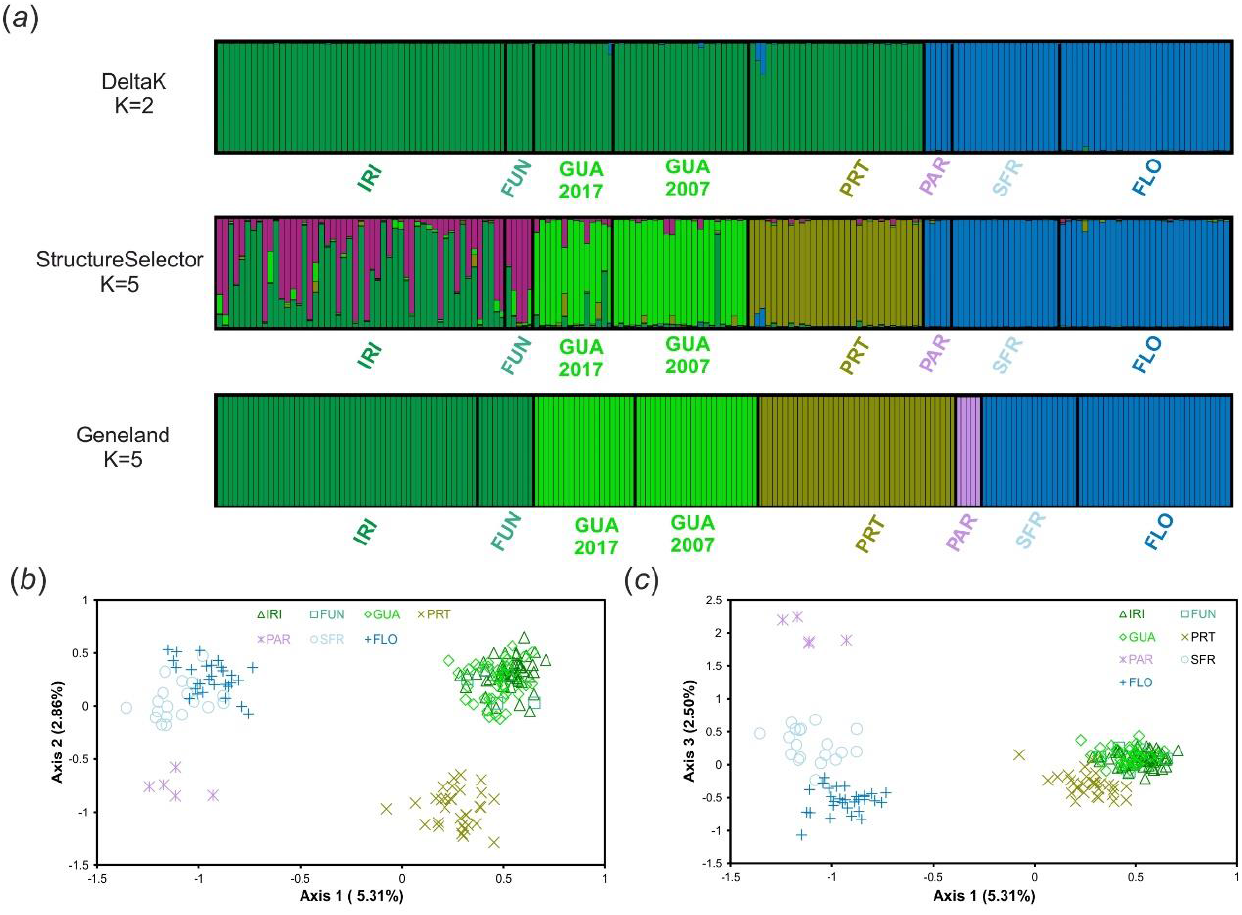
(*a*) Most likely genetic clusters (K) value for *Kryptolebias ocellatus* using 16 microsatellites analysed in Structure and Geneland. K values determined by ΔK method of Evanno et al. (2005) and metrics of Puechmaille (2016) implemented in STRUCTURESELECTOR. Geneland analysis includes spatial and mtDNA information, in addition to the microsatellite’s genotypes. Each individual is represented by a bar, and each colour represents a genetic cluster. (*b-c*) Factorial correspondence analysis for all *K. ocellatus* individuals coloured and shaped according to their sampling sites.

On average, 22.4 *K. ocellatus* individuals were genotyped at 16 microsatellite loci per sampling location, but sample size varied considerably (from 5 to 51) (Table 2). Overall, there was high level of variation at microsatellite loci in *K. ocellatus.* The number of alleles varied from 2 at locus R28 to 35 at locus R38, with an average of 17.6 alleles per locus considering all sampling locations combined. The mean expected heterozygosity (H_E_) was 0.56 (ranging from 0.47 to 0.60) for *K. ocellatus.* The *K. ocellatus* Northern clade populations (IRI, FUN and GUA) showed a higher average H_E_ (0.59) than Parati (0.53) and Southern clades (0.53). Only one sampling point (GUA dataset for 2007) had significant heterozygote deficiency (Table 2). Examination of single-locus F_IS_ values indicated that significant values of mean F_IS_ in were due to contribution of few loci and likely due to null alleles (Table S4). Since mean F_IS_ was non-significant after the atypical loci were excluded, we consider that all studied populations of *K. ocellatus* are in Hardy-Weinberg equilibrium and we kept all loci for further analyses (Table S4).

**Table 2.**
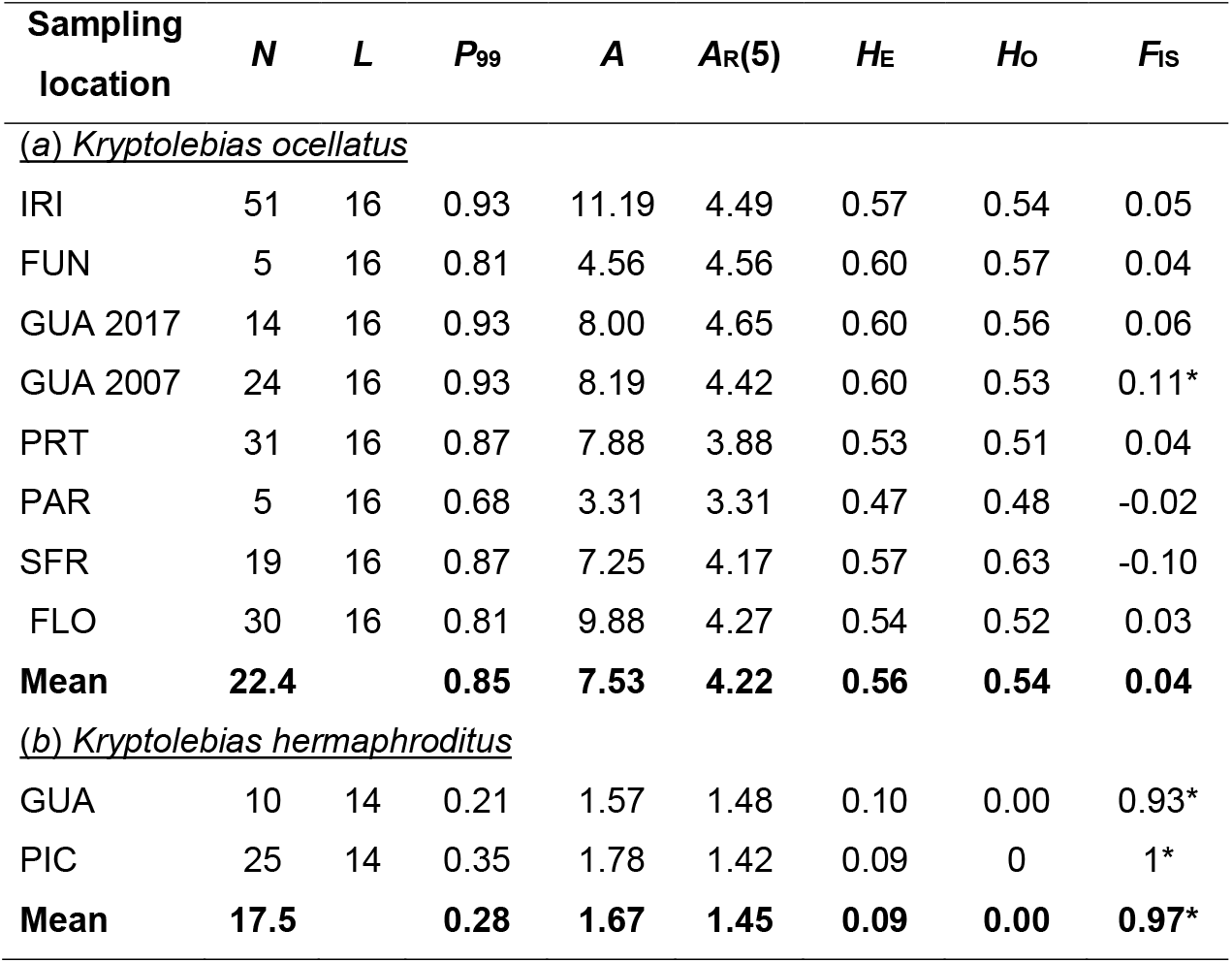
Descriptive statistics of genetic variation at microsatellite loci in (*a*) 179 *Kryptolebias ocellatus* (excluding putative hybrids) and (*b*) 35 *K. hermaphroditus* individuals. *Kryptolebias ocellatus* GUA samples from different years are separated. N = sample size; L = number of loci; P99 = proportion of polymorphic loci (99% criterion); A = average number of alleles; AR = allelic richness based on 5 individuals; H_E_ = expected heterozygosity; H_O_ = observed heterozygosity; F_IS_ = coefficient of inbreeding. Asterisks represent significant departures from HWE.

As expected for a selfing species, no loci were found to be under Hardy-Weinberg equilibrium for *K. hermaphroditus.* The number of alleles varied from 1 (at 50% of locus) to 10, with an average of 2.21 alleles per locus. No heterozygotes were found in both *K. hermaphroditus* populations. Significant heterozygote deficiency was detected for all loci that had more than one allele in *K. hermaphroditus* (Tables 2 and S4).

### Genetic differentiation and clustering analysis

Classification of individuals using STRUCTURE provided consistent results for each K across the 10 replicated runs. As expected in highly structured populations, the most divergent groups separate into distinct clusters first (Pritchard *et al*, 2000). Evanno’s ΔK method indicated the uppermost level of genetic structure was K = 2. This analysis indicated one genetic cluster encompassing fish from the Northern populations (IRI, FUN, GUA, PRT) and another composed of fish from the southernmost sampling sites (PAR, SFR and FLO) (Figs.1 and 3). Outcomes of K = 5 (indicated as the most likely number of genetic clusters by all metrics in STRUCTURESELECTOR; Fig. S3) assigned all fish from GUA and PRT to their own genetic clusters. Two genetic lineages were found to be admixed in the sampling points of Guanabara Bay (IRI and FUN). Southernmost sampling points were assigned to the same genetic cluster (Fig. 3). Geneland results incorporating mtDNA, microsatellites and spatial data generally agreed with those from STRUCTURE. Posterior distributions of the number of genetic clusters (K) showed a mode at K = 5 across all 10 replicated runs (Figs. 3, S3, and S4). Spatially, cluster 1 was composed of individuals from IRI and FUN while individuals from GUA, PRT and PAR each represented a unique genetic cluster (clusters 2, 3 and 4 respectively). Cluster 5 was composed of the southernmost individuals from SFR and FLO (Fig. S4). No differences between GUA samples from 2007 and 2017 were found across any clustering analysis. An additional Geneland analysis using 108 individuals with data for both mtDNA and microsatellites (excluding the hybrids), suggested K=4 as the most likely number of genetic clusters. Overall, the genetic clustering found in this analysis was similar to the one found using whole dataset (Fig. 3), with the exception that individuals from GUA have clustered with individuals from other populations of the Northern clade (IRI and FUN) (Fig. S5).

Factorial Correspondence Analysis (FCA) confirmed the uppermost subdivisions detected by Evanno’s method (ΔK) from STRUCTURE (Fig. 3). The plot along the two main axes showed that the major division was between the southern and the northern populations along axis 1. Along Axis 2, further genetic subdivision was found, matching Geneland results. PRT and PAR individuals formed separate clusters from Northern and Southern population, respectively. Along axis 3, PAR individuals differentiated even further (Fig. 3).

Genetic differentiation between populations of *K. ocellatus* was high and significant in global tests and pairwise comparisons. For example, the average F_ST_ among all pairwise comparisons was 0.25 (P < 0.001). In pairwise comparisons no difference was found between Guaratiba samples collected 10 years apart (F_ST_ = 0.00). F_ST_ in the remaining pairwise comparisons varied from 0.06 (between IRI and FUN; and SFR and FLO) to 0.38 (between IRI and PAR) (Table S6). The majority of pairwise F_ST_ were statistically significant after Bonferroni correction for multiple testing, with the exception of comparisons of FUN vs GUA-2017 (F_ST_ = 0.072), and FUN vs PAR (F_ST_ = 0.382), the latter most likely caused by small sample sizes in FUN and PAR (Table S5). Significant F_ST_ between PIC and GUA populations of *Kryptolebias hermaphroditus* was also found (0.30; P = 0.01).

Strong evidence for isolation by distance was found in *K. ocellatus* using both mtDNA (R^2^ = 0.58; P = 0.01) and microsatellites (R^2^ = 0.84; P = 0.003) pairwise genetic distances. In particular, two loci (R9 and R18) showed evident pattern of regional geographic differentiation between Northern and Southern populations, with little to no overlap in allele distribution between Northern and Southern populations (Fig. S6).

## Discussion

Theory predicts that, all else being equal, selfing should have magnified effects on genetic structure when compared to outcrossing species as a consequence of reduced effective population size and increased inbreeding (Charlesworth, 2003; Meunier *et al*, 2004). Here we found that, overall, *Kryptolebias ocellatus* populations across much of known species distribution are under Hardy-Weinberg equilibrium, strongly suggesting that despite androdioecious, *K. ocellatus* is mostly an outcrossing species. This finding supports early behavioural (Costa *et al*, 2010; Seegers, 1984), and genetic (Tatarenkov et al. 2009; for two populations only) indications of outcrossing as the main mating system in *K. ocellatus*, although the possibility that the species undergoes, even if only rarely, selfing cannot be fully discarded. It also remains to be established if hermaphrodites mate exclusively with males, or whether they can mate with each other. Our results also revealed deep population structuring in *K. ocellatus*, mostly following an pattern of isolation-by-distance (IBD), which generally contrasts with the high genetic homogeneity found in the morphologically-similar, predominantly-selfing and often-syntopic *K. hermaphroditus* across discontinuous mangrove forests along the Brazilian coast (Tatarenkov *et al*, 2009; 2011; 2017).

In the selfing species composing the *K. marmoratus* species complex, extensive genetic structure has been identified across (Tatarenkov *et al*, 2015; Tatarenkov *et al*, 2007), and within the same mangrove systems (Berbel-Filho *et al*, 2019; Ellison *et al*, 2012; Turko *et al*, 2018). An exception to this pattern of deep genetic structure is the high genetic homogeneity found among the selfing *K. hermaphroditus* populations across the Brazilian coast (Tatarenkov *et al*, 2011). As shown here, *K. hermaphroditus* from south-eastern Brazil carry a single *cox1* haplotype and are completely homozygous at polymorphic microsatellite loci. Geographically, this low genetic diversity scenario in *K. hermaphroditus* extends even further, with very little genetic differentiation in *K. hermaphroditus* populations from Southeast and Northeast Brazil, separated by approximately 2500 km along the coast (Tatarenkov *et al.* 2017). In contrast, in a much narrower geographic distribution (approximately 900 km along the coast from Magé in the State of Rio de Janeiro to Florianópolis in Santa Catarina state), *K. ocellatus* showed a deeper genetic structure with division in two genetic clusters (Northern and Southern), and moderate internal genetic structure within these clusters. In its relatively narrow geographic distribution, *K. ocellatus* also showed more *cox1* mtDNA haplotypes (22 vs 18) and higher interclade mtDNA genetic distances *(cox1* K2P distance: 1.1%) than the average genetic distance among clades *(cox1* K2P distance: 0.98% between ‘Central’ and ‘Southern’ clades) in the widely distributed (Florida (29°N) to São Paulo (23°S)) *K. marmoratus* species complex (Tatarenkov *et al*, 2017). Although the *K. ocellatus* phylogenetic reconstruction was based on a single mtDNA gene, which may not accurately represent the species tree, it was highly concordant with the microsatellite tree (Fig. S1), supporting the existence of at least two major clades across *K. ocellatus* distribution, with further genetic subdivisions within them. Thus, our results indicate that the two *Kryptolebias* species in south-eastern Brazil did not evolve by a sympatric speciation event in the region (Kanamori *et al*, 2016; Tatarenkov *et al*, 2017), and have remarkably different evolutionary history along the Brazilian coast, with *K. hermaphroditus* most likely being a recent coloniser of a mangrove area where *K. ocellatus* might have settled/originated much earlier.

Mangrove forests are typically associated to intertidal zones along rivers, estuaries and bay areas with brackish water (Ball, 1988; Hamilton and Casey, 2016). This association forms an overall discontinuous distribution of mangrove patches (Hamilton and Casey, 2016). Further contributing for mangrove forests fragmentation is human activity, which its effects are particularly pronounced in heavily urbanised areas, such as south-east Brazil (Branoff, 2017; Ferreira and Lacerda, 2016). The fragmented distribution of mangrove forests may have contributed for the pattern of IBD found here for *K. ocellatus,* with geographically more distant populations also being the most genetically dissimilar, both at mtDNA and microsatellites markers. The IBD pattern of genetic structure has also been found for highly selfing populations of *K. marmoratus* in Florida (Tatarenkov et al. 2015), indicating that, in some occasions (but see below), long-distance dispersal in mangrove killifishes is limited. Mangrove killifish are the only rivulid species living in brackish waters (Costa *et al*, 2010) and rarely share mangrove microhabitats with other fish species permanently (Taylor, 2012). Therefore, making inferences between the genetic structure of mangrove killifishes and other mangrove-dwelling fish species is challenging. Studies of various mangrove tree species showed weak genetic structure among estuaries in south-eastern Brazil, with a general north-south pattern of dispersal, guided by the Brazilian ocean current (Francisco *et al*, 2018; Mori *et al*, 2015; Pil *et al*, 2011). This high gene flow scenario among different estuaries has also been observed in other mangrove-dwelling species in the same region which disperse through pelagic larvae, such as crabs (Britto *et al*, 2018; de Oliveira-Neto *et al*, 2008; Oliveira-Neto *et al*, 2007). The strong genetic subdivision found in *K. ocellatus* between Northern and Southern estuaries in southwestern Atlantic (with particularly high F_ST_ values at the mtDNA), contrasts with the general pattern of panmixia pattern mentioned above for mangrove-associated species in the same region. Although our data indicates that *K. ocellatus* reproduces mostly via outcrossing, we cannot discard that the high genetic differentiation between Northern and Southern populations could have been amplified by geographical variation on ancestral events of selfing. In addition, given the strong indication of IBD, the high differentiation between Northern and Southern populations may have been magnified by the lack of sampling in more intermediate locations (e.g. along São Paulo state coast, Fig. 1). Finally, further research is needed to indicate whether hybridisation (and potential ancestral introgression) between *K. ocellatus* and *K. hermaphroditus* may have influenced the allele distribution and population differentiation of *K. ocellatus* in the Northern populations.

The deep genetic structure and limited dispersal between estuaries of *K. ocellatus* also contrasts with the long-distance dispersal capacity observed in *K. hermaphroditus* along the Brazilian coast (Tatarenkov *et al*, 2017) could be due to differences in colonisation success as a result of their different mating systems. Previous research in plants indicated that selfing is associated with increased dispersal capacity and colonisation success (de Waal *et al*, 2014). Mangrove killifishes are poor swimmers, but their long-term dispersal mangrove killifishes can be facilitated by adhesive eggs transported via floating material (Tatarenkov *et al*, 2012; Turko and Wright, 2015). In *K. hermaphroditus,* self-fertilisation provides the possibility for a single individual to found a new population after a long-distance dispersal event, while *K. ocellatus* would require at least two individuals to breed. This hypothesis is supported by the large combined geographic range of the selfing mangrove killifishes (*K. marmoratus* and *K. hermaphroditus*), extending from Florida (23°N) to south Brazil (29°S), although further research is needed to investigate how the differences in mating systems between *K. ocellatus* and *K. hermaphroditus* can influence their colonisation capacities.

## Conclusions

Contrary to the theorical predictions that selfing species should result in high population structuring given its reduced effective population size due to inbreeding, we found that the outcrossing species *K. ocellatus* had stronger population structure in a narrower geographical range than its morphologically-similar and often syntopic selfing species *K. hermaphroditus*. These findings highlight that other factors such as colonisation time, extent of gene flow, dispersal and colonisation success may have more profound effects on the current patterns of population structure than differences in mating systems between selfing and outcrossing species.

## Acknowledgements

We are grateful to ICMbio for providing help with accommodation and facilities, especially the teams working at Parque Estadual Serra do Mar: Núcleo Picinguaba and Parque Estadual Serra do Mar: Estação Ecológica Juréia-Itatins. We are thankful to Dr. Ingo Schlupp from Oklahoma University for his friendly review. We also thank to Dr. Joana Robalo and two other anonymous reviewers whose comments and suggestions substantially improved the manuscript.

## Ethics

This work followed the Swansea ethics committee guidelines. Sampling work was carried out under license ICMBio/SISBIO 57145-1/2017

## Author’s contribution

SC, WMB-F, AT, SMQL and CGL conceived the study and obtained the funding. WMB-F, HMVES, ML, SMQL collected the samples. WMB-F and AT carried out the genetic analyses. WMB-F wrote the manuscript with participation of all authors

## Supporting information

### Taxonomic nomenclature of *Kryptolebias* species from southeast Brazil

The taxonomic status of mangrove killifishes species from in southeast Brazil has gone through several rearrangements, being particulaly complicated by historical poor species diagnosis and morphological similarities among sympatric species (Costa, 2011; Costa, 2016; Seegers, 1984). *K. ocellatus* was intially described by Hensel (1868) using a single specimen from Rio de Janeiro, Brazil. Seegers (1984) suggested the existence two syntopic species in Rio de Janeiro, the hermaphroditic *K. ocellatus*, and a yet undescribed dimorphic species, which was named as *K. caudomarginatus*. Two decades later, Costa (2011) argued that the species originally described by Hensel as *K. ocellatus,* actually referred to a species with hermaprhodites and males, reclassifying *K. caudomarginatus* as a junior synonym of *K. ocellatus,* while the other species composed only of selfing hermaphrodites, was named *K. hermaphroditus* (Costa, 2011). However, the discussion about the taxonomic status of these mangrove killifish species is still undergoing, and further taxonomic rearrangements may happen with increasing taxonomical research (Huber, 2017; Tatarenkov *et al*, 2020; Tatarenkov *et al*, 2017).

Before the taxonomic review by Costa (2011), the current *K. ocellatus* (composed by males and hermaphrodites in approximately similar sex ratio, (Costa, 2016)) was called *K. caudomarginatus* (Seegers, 1984) while the selfing species (the current *K. hermaphroditus)* was referred to as *K. ocellatus.* Therefore, in the previous studies from where we used part of the data analysed here (Tatarenkov *et al*, 2011; Tatarenkov *et al*, 2009), *K. ocellatus* is referred to as *K. caudomarginatus* and *K. hermaphroditus* as *K. ocellatus*.

**Figure S1.**
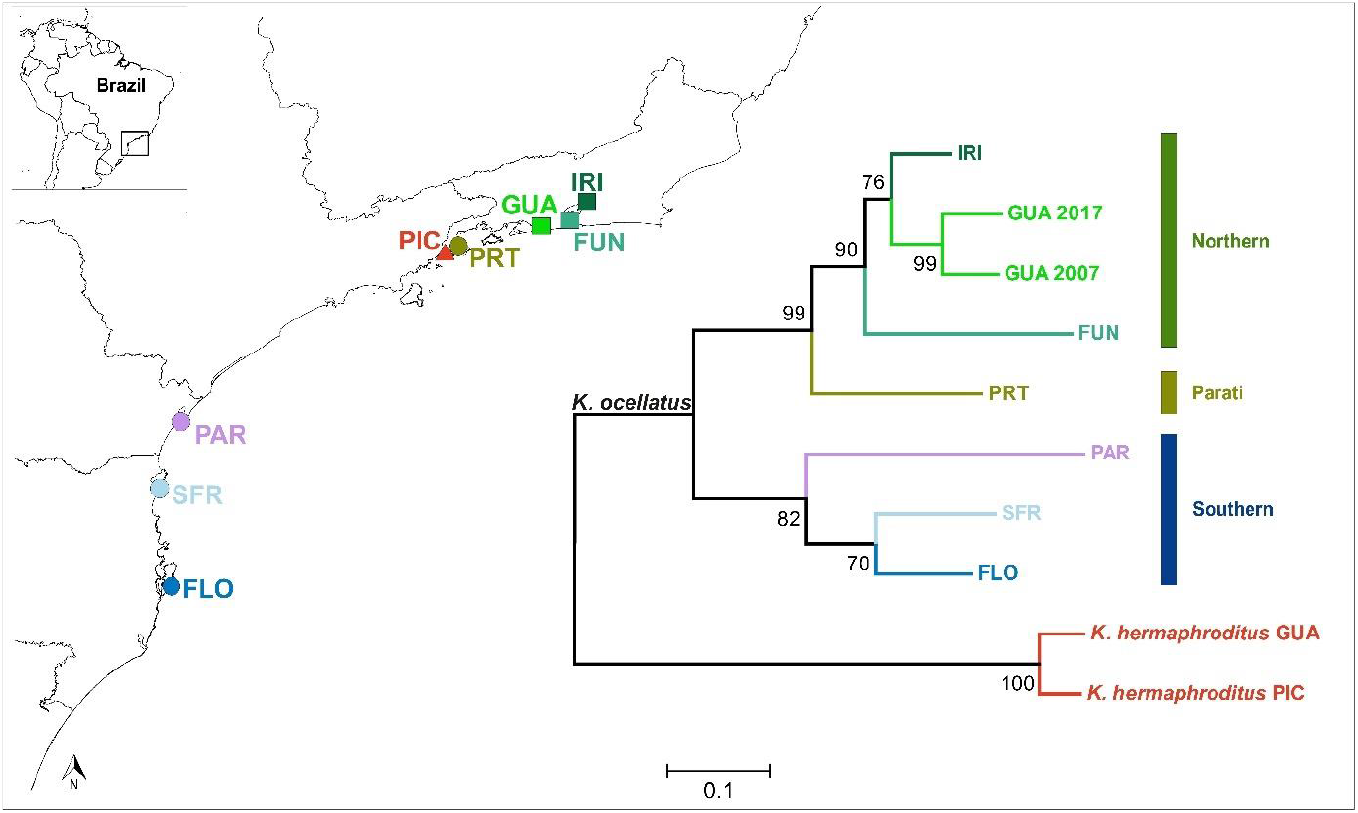
Neighbour-Joining tree for populations of *K. ocellatus* (excluding the hybrids) and *K. hermaphroditus.* Nei’s (1978) genetic distances were estimated from allele frequencies at 16 microsatellites loci. Colours represent different sampling locations described in Table 1. Bars represent major mtDNA clades. Numbers on nodes represent bootstrap support after 1000 replications.

**Figure S2.**
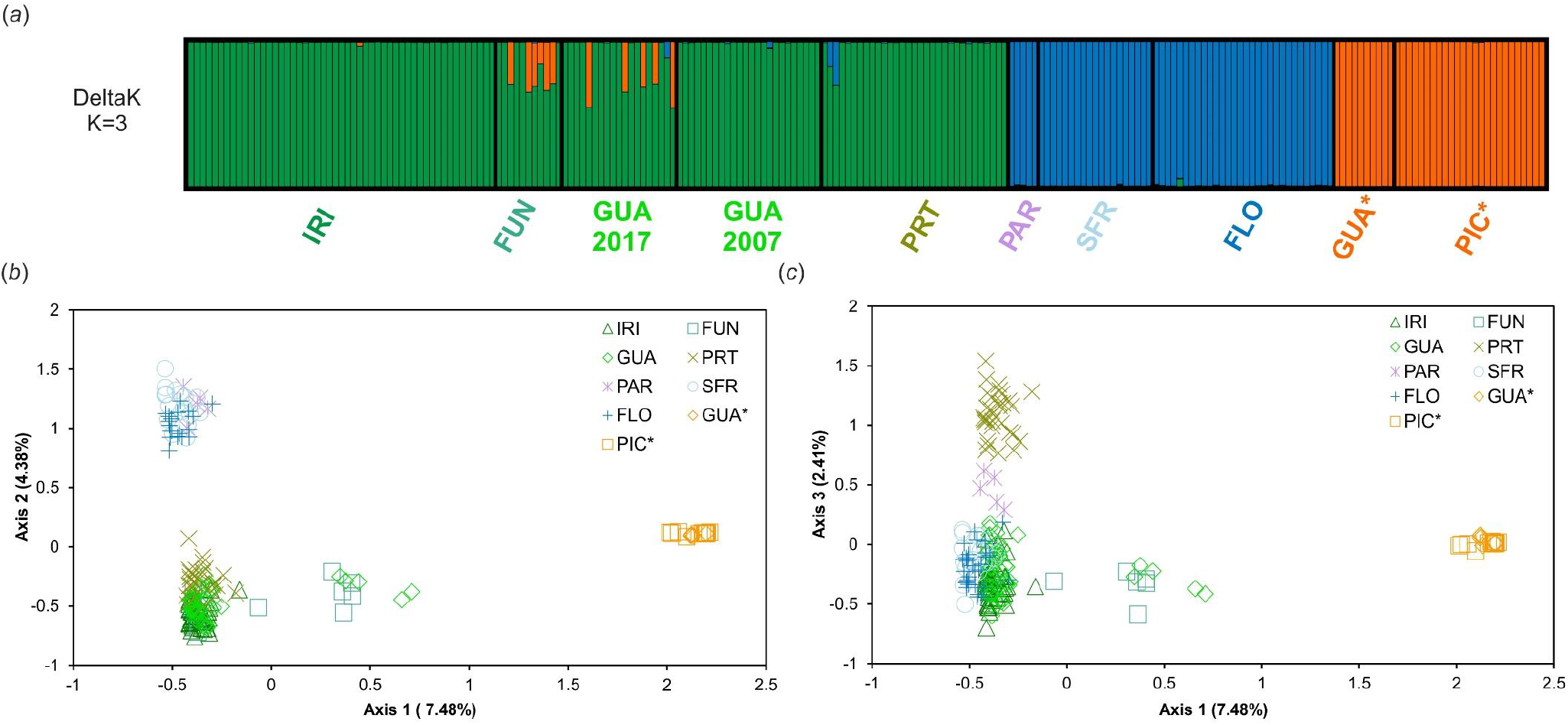
(*a*) Panel showing the most likely genetic clusters (K) value for the 16 microsatellites genotyped in 190 *Kryptolebias ocellatus* (including potential hybrids) and *K. hermaphroditus* (from Tatarenkov et al. 2011) ran in Structure and determined by ΔK method of Evanno et al. (2005). Each individual is represented by a bar, and each colour represents a genetic cluster. *(b-c)* Factorial correspondence analysis for all *K. ocellatus* individuals coloured and shaped according to their sampling sites. Sampling points are described in Table 1. Asterisks represent sampling points for *K. hermaphroditus*.

**Figure S3.**
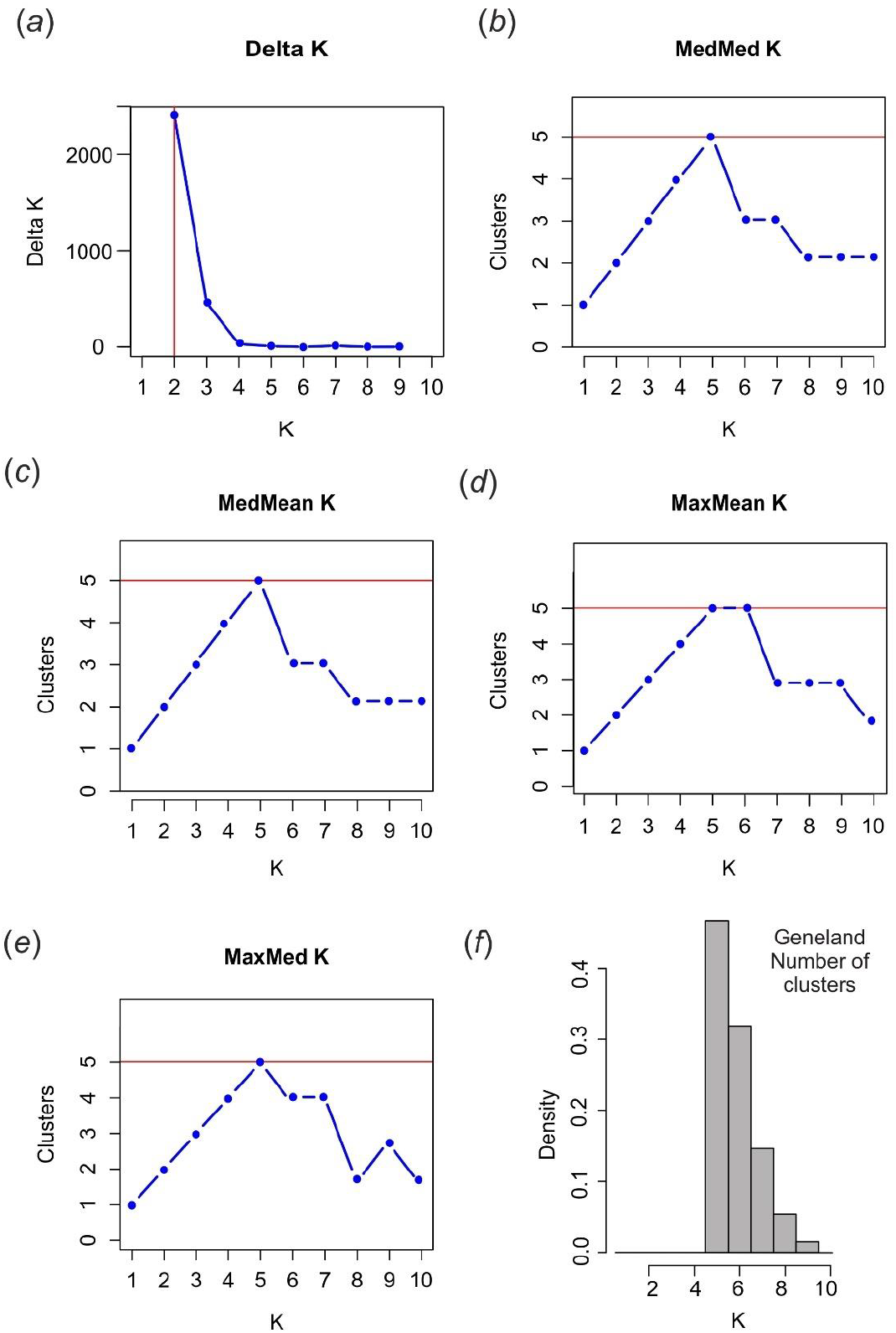
Estimated number of genetic clusters (K) from different methods. (*a*) deltaK method of Evanno *et al* (2005); (*b*) median of medians (MedMedK), (*c*) medians of means (MedMeanK); (*d*) maximum of medians (MaxMedK) and (*e*) maximum of the means (MaxMeaK) as implemented by Puechmaille (2016) to account for unevenness of sampling sizes and hierarchical structure; (*f*) posterior density distribution of the number of clusters estimated from Geneland.

**Figure S4.**
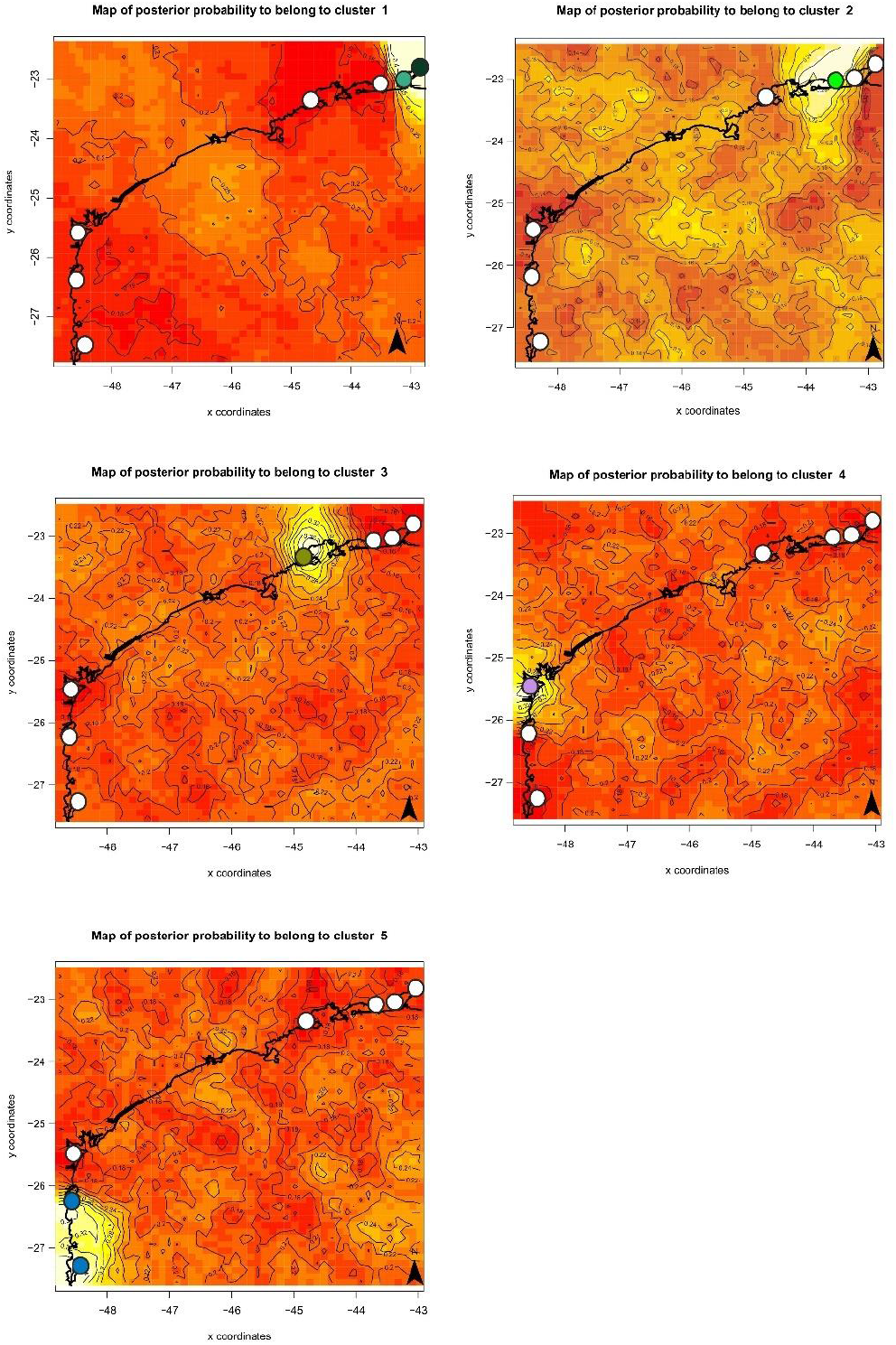
Maps of Geneland individual assignments to clusters for K = 5 with geographical coordinates. The highest membership values are in light yellow at the background.

**Figure S5.**
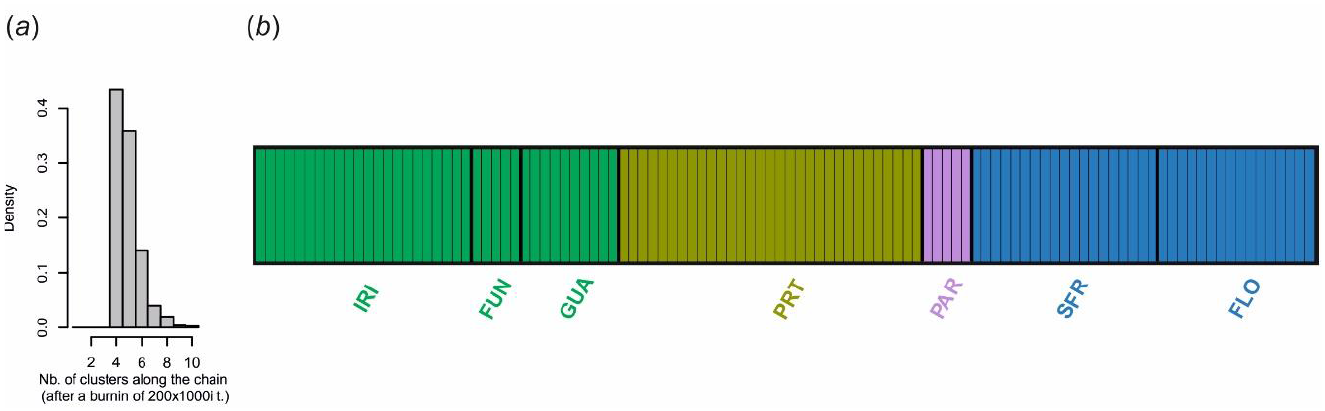
Geneland analysis results using only individuals which contained both *cox1* sequence and microsatellite genotypes (108 individuals excluding hybrids). (*a*) Posterior density distribution of the number of clusters estimated from Geneland; (*b*) admixture plot for all individuals for K= 4 as indicated as the most likely number of genetic clusters in the dataset.

**Figure S6.**
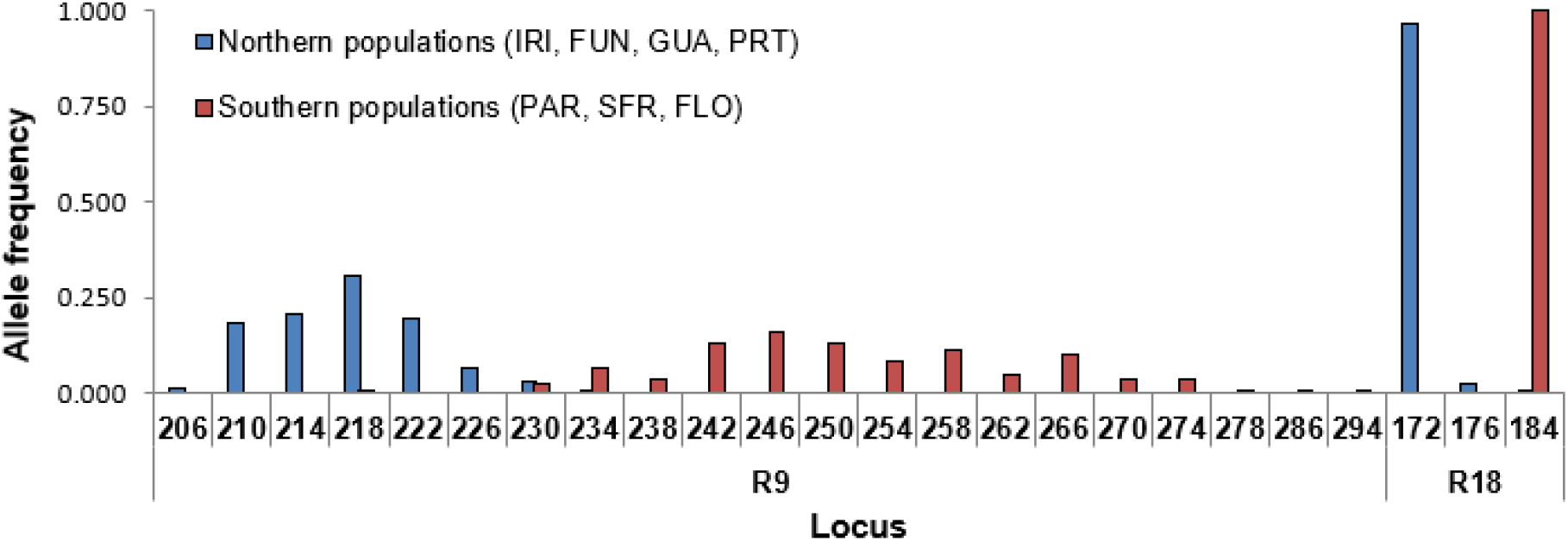
Allele frequency for locus R9 and R18 between Northern and Southern populations of *Kryptolebias ocellatus*.

**Table S1.** *cox1* haplotype distribution across different locations found for *Kryptolebias ocellatus* and *K. hermaphroditus.*

|  | *K. ocellatus* |  |  |  |  |  |  |  |  |  |  |  |  |  |  |  |  |  |  |  |  |  | *K.*<br>*hermaphroditus* |
| --- | --- | --- | --- | --- | --- | --- | --- | --- | --- | --- | --- | --- | --- | --- | --- | --- | --- | --- | --- | --- | --- | --- | --- |
| Haplotype<br>number | 1 | 2 | 3 | 4 | 5 | 6 | 7 | 8 | 9 | 10 | 11 | 12 | 13 | 14 | 15 | 16 | 17 | 18 | 19 | 20 | 21 | 22 | 1 (Kherm) |
| 1. IRI | 17 | 1 | 2 | 1 | 1 |  |  |  |  |  |  |  |  |  |  |  |  |  |  |  |  |  |  |
| 2. FUN | 1 |  |  | 3 |  | 6 | 1 |  |  |  |  |  |  |  |  |  |  |  |  |  |  |  | 29 |
| 3. GUA |  |  |  |  |  |  |  | 15 | 2 |  |  |  |  |  |  |  |  |  |  |  |  |  | 30 |
| 4. PRT |  |  |  |  |  |  |  |  |  | 6 | 1 | 2 | 30 |  |  |  |  |  |  |  |  |  |  |
| 5. PAR |  |  |  |  |  |  |  |  |  | 1 |  |  |  | 1 | 3 |  |  |  |  |  |  |  |  |
| 6. SFR |  |  |  |  |  |  |  |  |  |  |  |  |  | 1 | 4 | 5 | 1 | 1 | 6 | 1 |  |  |  |
| 7. FLO |  |  |  |  |  |  |  |  |  |  |  |  |  | 11 |  |  |  |  |  |  | 3 | 2 |  |
| 8. PIC |  |  |  |  |  |  |  |  |  |  |  |  |  |  |  |  |  |  |  |  |  |  | 2 |
| **Total** | **18** | **1** | **2** | **4** | **1** | **6** | **1** | **15** | **2** | **7** | **1** | **2** | **30** | **13** | **7** | **5** | **1** | **1** | **6** | **1** | **3** | **2** | **61** |

**Table S2.** Descriptive statistics of genetic variation at *cox1* mitochondrial DNA (mtDNA) gene for clades and sampling locations for (*a*) *K. ocellatus* and (*b*) *K. hermaphroditus.* H = number of haplotypes; S = number of polymorphic sites; *h* = haplotype diversity; *π* = nucleotide diversity.

|  | Population diversity indices |  |  |  |  |
| --- | --- | --- | --- | --- | --- |
| | *N* | *H* | *S* | *h* | $\pi$ |
| **(a) *K. ocellatus*** | 129 | 22 | 26 | 0.895 | 0.007 |
| *Major mtDNA clades* |  |  |  |  |  |
| Northern clade | 50 | 9 | 10 | 0.775 | 0.002 |
| Parati | 39 | 4 | 3 | 0.391 | 0.0007 |
| Southern clade | 40 | 10 | 15 | 0.836 | 0.002 |
| *Sampling locations* |  |  |  |  |  |
| IRI | 22 | 5 | 5 | 0.407 | 0.001 |
| FUN | 11 | 4 | 6 | 0.673 | 0.002 |
| GUA | 17 | 2 | 1 | 0.221 | 0.0004 |
| PRT | 39 | 4 | 3 | 0.391 | 0.0007 |
| PAR | 5 | 3 | 9 | 0.700 | 0.006 |
| SFR | 19 | 7 | 4 | 0.819 | 0.002 |
| FLO | 16 | 3 | 3 | 0.508 | 0.002 |
| **(b) *K. hermaphroditus*** | 61 | 1 | 0 | 0 | 0 |

**Table S3.** Descriptive statistics of genetic variation at *cox1* mitochondrial DNA (mtDNA) gene for clades and sampling locations for *K. ocellatus* excluding mtDNA sequences from 11 hybrids individuals (six from FUN and five from GUA). H = number of haplotypes; S = number of polymorphic sites; *h* = haplotype diversity; *π* = nucleotide diversity.

|  | Population diversity indices |  |  |  |  |
| --- | --- | --- | --- | --- | --- |
| | *N* | *H* | *S* | *h* | $\pi$ |
| ***K. ocellatus*** | 118 | 21 | 23 | 0.886 | 0.007 |
| *Major mtDNA clades* |  |  |  |  |  |
| Northern clade | 39 | 8 | 7 | 0.725 | 0.003 |
| Parati | 39 | 4 | 3 | 0.391 | 0.0007 |
| Southern clade | 40 | 10 | 15 | 0.836 | 0.002 |
| *Sampling locations* |  |  |  |  |  |
| IRI | 22 | 5 | 5 | 0.407 | 0.001 |
| FUN | 5 | 3 | 3 | 0.800 | 0.002 |
| GUA | 12 | 2 | 1 | 0.303 | 0.0004 |
| PRT | 39 | 4 | 3 | 0.391 | 0.0007 |
| PAR | 5 | 3 | 9 | 0.700 | 0.006 |
| SFR | 19 | 7 | 4 | 0.819 | 0.002 |
| FLO | 16 | 3 | 3 | 0.508 | 0.002 |

**Table S4.** Pairwise F_ST_ values (below diagonal), Kimura-2-parameter (K2P) genetic distance (in percentage above diagonal) and within group K2P distance (in the diagonal) for mtDNA 129 *cox1* gene among sampling locations for *Kryptolebias ocellatus. Asterisks represent p-value ≤ 0.05.*

|  | **1** | **2** | **3** | **4** | **5** | **6** | **7** |
| --- | --- | --- | --- | --- | --- | --- | --- |
| 1. IRI | **0.1** | 0.4 | 0.2 | 0.9 | 1.1 | 1.2 | 1.1 |
| 2. FUN | 0.55* | **0.2** | 0.3 | 1.0 | 1.2 | 1.3 | 1.2 |
| 3. GUA | 0.58* | 0.59* | **0** | 0.7 | 0.9 | 1.1 | 0.9 |
| 4. PRT | 0.88* | 0.91* | 0.88* | **0.1** | 1.1 | 1.4 | 1.3 |
| 5. PAR | 0.77* | 0.79* | 0.69* | 0.85* | **0.6** | 0.5 | 0.4 |
| 6. SFR | 0.83* | 0.84* | 0.80* | 0.90* | 0.14 | **0.3** | 0.3 |
| 7. FLO | 0.87* | 0.90* | 0.85* | 0.92* | 0.28* | 0.30* | **0.1** |

**Table S5.** Pairwise F_ST_ values (below diagonal), Kimura-2-parameter (K2P) genetic distance (in percentage above diagonal) and within group K2P distance (in the diagonal) for mtDNA 118 *cox1* gene among sampling locations for *Kryptolebias ocellatus (excluding 11 hybrid individuals). Asterisks represent p-value ≤ 0.05.*

|  | **1** | **2** | **3** | **4** | **5** | **6** | **7** |
| --- | --- | --- | --- | --- | --- | --- | --- |
| 1. IRI | **0.1** | 0.3 | 0.2 | 0.9 | 1.1 | 1.2 | 1.1 |
| 2. FUN | 0.50* | **0.2** | 0.3 | 0.9 | 1.1 | 1.3 | 1.1 |
| 3. GUA | 0.53* | 0.56* | **0** | 0.7 | 0.9 | 1.1 | 0.9 |
| 4. PRT | 0.88* | 0.89* | 0.90* | **0.1** | 1.1 | 1.4 | 1.3 |
| 5. PAR | 0.77* | 0.62* | 0.76* | 0.85* | **0.6** | 0.5 | 0.4 |
| 6. SFR | 0.83* | 0.79* | 0.83* | 0.90* | 0.14 | **0.3** | 0.3 |
| 7. FLO | 0.87* | 0.86* | 0.89* | 0.92* | 0.28* | 0.30* | **0.1** |

**Table S6.** F_IS_ values per locus and population for 179 individuals of *Kryptolebias ocellatus* and 35 *K. hermaphroditus.* Asterisks indicate significance (p<0.05) after correction for multiple testing. Hybrids excluded from analysis. GUA samples from different years are separated.

|  | *K. ocellatus* |  |  |  |  |  |  |  | *K. hermaphroditus* |  |
| --- | --- | --- | --- | --- | --- | --- | --- | --- | --- | --- |
|  | IRI | FUN | GUA 2017 | GUA 2007 | PRT | PAR | SFR | FLO | GUA | PIC |
| **R9** | -0.05 | 0.48 | 0.33* | 0.59* | 0.05 | -0.17 | -0.12 | 0.07 | NA | NA |
| **R11** | 0.09 | -0.08 | 0.06 | 0.13 | 0.25* | -0.06 | -0.10 | -0.02 | NA | NA |
| **R18** | -0.01 | 0.00 | -0.04 | 0.00 | -0.01 | NA | NA | NA | 1.00* | NA |
| **R23** | 0.01 | -0.05 | -0.04 | 0.05 | 0.02 | 0.29 | -0.04 | 0.17 | NA | 1.00* |
| **R27** | 0.03 | 0.17 | -0.06 | -0.00 | -0.02 | 0.05 | -0.06 | 0.04 | 1.00* | NA |
| **R28** | NA | NA | NA | NA | NA | NA | 0.000 | NA | NA | NA |
| **R30** | 0.26* | 0.20 | -0.15 | -0.02 | -0.13 | 0.03 | -0.05 | -0.01 | NA | 1.00 |
| **R33** | 0.05 | 0.05 | 0.05 | 0.00 | 0.07 | -0.20 | -0.18 | NA | NA | 1.00* |
| **R34** | 0.00 | NA | 0.00 | -0.06 | -0.01 | NA | -0.71* | -0.09 | NA | NA |
| **R37** | -0.01 | -0.08 | 0.11 | 0.04 | 0.01 | -0.14 | 0.00 | 0.05 | 0.88* | 1.00* |
| **R38** | 0.06 | -0.08 | 0.26* | 0.16 | 0.03 | -0.11 | -0.08 | 0.16 | NA | 1.00* |
| **B86** | -0.09 | 0.27 | -0.13 | 0.07 | -0.19 | -0.33 | -0.05 | -0.18 | NA | ANA |
| **R90** | -0.01 | NA | -0.02 | -0.07 | 0.17 | 0.17 | -0.32 | -0.10 | NA | NA |
| **R92** | 0.42* | 0.04 | 0.37 | 0.23 | 0.05 | 0.00 | 0.25 | -0.05 | NA | NA |
| **R103** | -0.11 | -0.14 | -0.06 | 0.21 | -0.05 | NA | -0.04 | 0.02 | NA | NA |
| **R112** | -0.14 | -0.28 | -0.23 | 0.06 | NA | NA | NA | 0.00 | NA | NA |
| **All** | **0.05\*** | **0.04** | **0.06\*** | **0.11\*** | **0.04** | **-0.02** | **-0.10\*** | **0.03** | **0.93\*** | **1.00\*** |

**Table S7.**
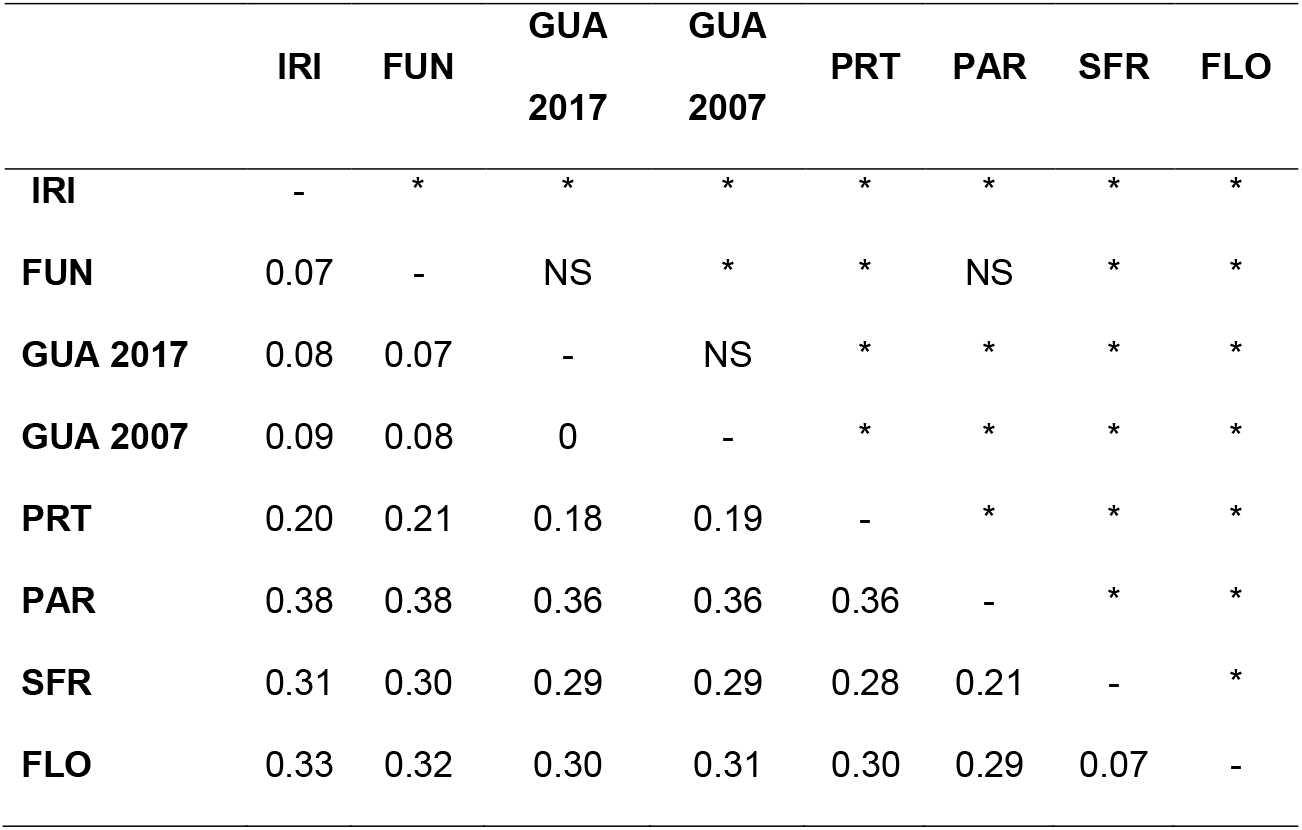
Pairwise F_ST_ values among *Kryptolebias ocellatus* populations based on 16 microsatellite loci. Potential hybrids are excluded from the analysis. Asterisks indicate significance (p<0.05) after Bonferroni correction. GUA samples from different years are separated.

|  | IRI | FUN | GUA<br>2017 | GUA<br>2007 | PRT | PAR | SFR | FLO |
| --- | --- | --- | --- | --- | --- | --- | --- | --- |
| **IRI** | - | * | * | * | * | * | * | * |
| **FUN** | 0.07 | - | NS | * | * | NS | * | * |
| **GUA 2017** | 0.08 | 0.07 | - | NS | * | * | * | * |
| **GUA 2007** | 0.09 | 0.08 | 0 | - | * | * | * | * |
| **PRT** | 0.20 | 0.21 | 0.18 | 0.19 | - | * | * | * |
| **PAR** | 0.38 | 0.38 | 0.36 | 0.36 | 0.36 | - | * | * |
| **SFR** | 0.31 | 0.30 | 0.29 | 0.29 | 0.28 | 0.21 | - | * |
| **FLO** | 0.33 | 0.32 | 0.30 | 0.31 | 0.30 | 0.29 | 0.07 | - |

